# The male-biased sex ratio in humans and its role in the transition from promiscuity to pair bonding

**DOI:** 10.64898/2025.12.01.691708

**Authors:** Matthew C. Nitschke, Kristen Hawkes, Peter S. Kim

## Abstract

Despite decades of interest and multiple hypotheses formulated to explain the evolution of monogamy, there remains considerable disagreement about its frequency among our primate relatives and its origins in the human lineage. Two recent reviews point out inconsistencies in both the terminology used to categorise observed behaviour and the robustness of data sets commonly used to justify these hypotheses, underscoring the importance of careful analysis. Agent-based mathematical models can help mitigate some of these difficulties by providing simplified models that allow us to focus on important details and add precision to the processes involved. Here, we introduce such a model that permits us to include key life history differences between chimpanzees and humans including our unique longevity, length of post-menopausal lifestage, shorter birth intervals, and the length of our preferential relationships. To do this, we include explicit age structure, controlling for the length of the interbirth interval and mean adult longevity. By adjusting for these two parameters, we determine the dominant strategy by which males compete for paternities. We show that this change aligns with a shift in the Operational Sex Ratio (OSR), which measures the degree of competition required for each available female. We further investigate the dynamics resulting from both paternity uncertainly– caused by multiple mating males obtaining a paternity from a guarded female–and pair-bond breakup. Although several factors likely played a role in the transition to monogamy in humans, our results indicate that a male-biased sex ratio and the subsequent increase in competition for additional paternities was among them.

## 1 Introduction

Human life history patterns have diverged from those of other hominids in many important ways [8, 34]. Humans grow more slowly and have an extended period of juvenile dependence, produce first offspring at a later age, and enjoy a longer lifespan [1, 13, 15, 16, 36].

Despite this, humans maintain short birth intervals compared to other hominids. Most importantly, lifespans are longer than any other primate, yet in humans, female fertility ceases well before the end of average adult lifespans. Although age at menopause is about the same in chimpanzees and humans [46], chimpanzees age faster, rarely outliving their fertility. In contrast, women can expect substantial postmenopausal survival [26]. These differences undoubtedly contributed to human mating behaviour that is also unique among hominids. Unlike our closest relatives, it is only humans who form long-term preferential mating relationships. However, the transition from ancestral promiscuity to pair bonding and monogamous behaviour continues to be a puzzle.

Multiple studies have sought to explain this transition [6, 30, 38], yet no consensus has been reached on the underlying mechanisms (see [37] for a brief overview of the leading theories). Opie *et al*., using comparative phylogenetic analyses, argued that infanticide prevention was the principal force driving monogamy in our lineage [38]. In contrast, that same year Lukas and Clutton-Brock employed similar methods to conclude that male infanticide alone could not account for the evolution of social monogamy across mammals; instead, they pointed to female–female intolerance and low female density—which constrain males’ ability to monopolise multiple mates—as more likely drivers [30]. These divergent findings highlight the complexity of this evolutionary transition and the ongoing debate it generates. As discussed in a recent review [17], comparative analyses can offer valuable insights but must be applied with great caution when combining different data sets.

Although numerous hypotheses have been proposed to explain the origins of monogamy in mammals, it almost certainly emerged through co-evolutionary processes rather than any single driver [17, 25]. Many investigations have illuminated the mechanisms underpinning monogamous behaviour, yet few account for distinctive human life-history traits, most notably the male-biased adult sex ratio and support from post-menopausal females. Using mathematical models, Gavrilets [6] showed that between-individual variation is crucial to understanding the social dilemmas [11] and behaviours that contribute to pair bonding in humans. However, he used assumptions inconsistent with ethnographic data; both the Aché and Hadza showed men target prey that everyone consumes with no differential benefits to their own wives and children [14, 20]. Here, we incorporate some of this variation with a simple model that includes important life-history traits and focuses on a mechanistic understanding of how mate-guarding emerged in our lineage. With a male-biased sex ratio in the fertile ages, the competing pool of males makes claiming a female and defending that claim the strategy that wins more paternities rather than leaving to compete for paternity opportunities elsewhere [43]. Our objective here is to investigate the factors that led to the earliest appearance of mate guarding in our own lineage, a shift from the ancestral multiple mating strategy to persistent pair bonding.

Several recent mathematical models investigate the mechanisms behind this hypothesis by exploring the links between mating strategies, sex ratios and partner availability [27,28,40, 43,44]. Two such ratios appear frequently in the literature, the adult sex ratio (ASR), defined as the ratio of males to females in the fertile ages [43], and the operational sex ratio (OSR), which counts only the subset of adults currently capable of conception [4]. Each ratio captures a different aspect of male competition; higher OSR corresponds to more competitions for each paternity opportunity, and a higher ASR to more competitions for each fertile female.

One of the more recent such models was provided by Schacht and Bell [43] who include three male strategies: mate-guarding, multiple-mating and parental care. Under broad assumptions, they show the large effect of the ASR on male strategies concluding that mate guarding, rather than paternal care, likely propelled the evolution of human monogamy. Loo *et al*. [27, 28] significantly extended this work by allowing for guarding inefficiency and pair-bond break-up. They construct a system of ordinary differential equations that includes an investigation into the effects of changes to parameters on the long-term equilibria of these strategies. The work of both Schacht and Bell [43] and Loo *et al*. [27, 28] together support the hypothesis that mate-guarding is one of the primary forces behind the evolution of pair bonding. However, these models only include populations of males. A more recent model builds on these results by including additional populations of females and offspring to explore the link between the ASR and dominant male strategies [40]. They include populations of females and subsequent offspring associated with males employing a particular strategy.

We build on our previous model [37] to develop a probabilistic, agent-based model of male mating strategies. This permits us to add age structure to the model that was previously limited. We focus our attention on the chimpanzee-human relationship and concentrate on two strategies: mate-guarding and multiple-mating. This approach allows us to simplify the dynamics and focus on the two principal parameters affecting the viability of a given strategy: the length of the interbirth interval and mean adult longevity. A key part of this story is that females are not fertile at postmenopausal ages while males keep making new sperm each day during those same years, which significantly increases the number of males in the mating pool. We show that the emergence of guarding as the dominant mating strategy is most sensitive to an increase in expected adult longevity, since more males competing over a smaller selection of fertile females increases the competition for mates. However, the prevailing strategy is also affected by other important parameters in less ideal cases such as the average pair bond length and paternity uncertainty.

## 2 Model

We consider an agent-based model (ABM) in which males either guard, G, or multiply mate, M. The model classifies individuals into stages based on age, dependency state, sex, mating strategy, and child-bearing status. Thus, each individual advances through several life stages. We further assume a framework in which mature females are either *in* and available to mate, *out* and unable to do so because they are providing care, or are postmenopausal. Guarding males are faithful to their partners and thus remain out of the competition for additional females while they are in a committed pair. Multiple-mating males return to the mating pool after each successful mating or unsuccessful attempt. See Table S1 in the Supplementary Information for a full list of populations we consider.

An overview of the timeline for individual life stages is illustrated in Figure 1. All individuals go through distinct stages as they age including infant dependence (both mother and infant are grouped together in this stage), juvenile independence, sexual maturity, post-fertility and death. Note that once infants are weaned, mothers are available for conception again. At this stage, chimpanzee youngsters are fully independent; however, humans are still mostly dependent. To simplify the dynamics, we do not include this distinction in our model directly. Instead, we approximate these differences by adjusting the fixed interbirth interval [10,22,23]. Thus, humans owe a shorter interbirth interval to post-fertile females. Females experience an end to fertility due to menopause. Males leave the mating pool due to frailty. The specific features of the ABM are detailed in the subsections below.

**Figure 1.**
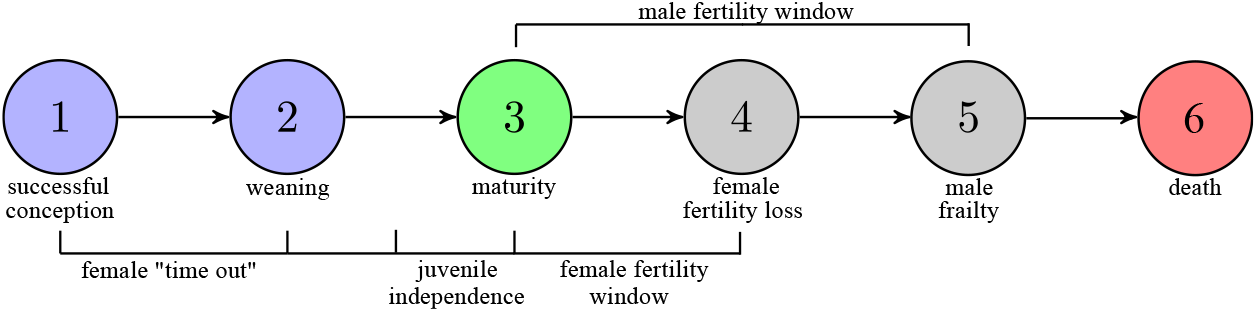
Lifetime events. Individuals go through a sequence of events throughout their life-time. (1) Successful conception where females temporarily leave the mating pool (time out) to care for dependants. (2) Infants are weaned and mothers can rejoin the mating pool. Juveniles at this stage are assigned a sex and mating strategy. Note that there is a period between independence from mothers and full juvenile independence where human youngsters still need help from adults. For simplicity, we do not consider the dynamics of parental care and consider all juveniles independent once weaned. (3) Juveniles become sexually mature and enter the mating pool at a set age. This is the point where they become adults and stay until their fertility ends or they die. (4) The fertility window ends for females at age 45. Paired males return to the mating pool to compete for other available females. (5) Males leave the active mating pool due to frailty. (6) All members of the population die or are no longer tracked after age 80.

### 2.1 Infant dependency, juvenile independence, sexual maturity and death

Individuals advance through a period of infant dependency, juvenile independence and sexual maturity. Once a male and female have a successful conception, the female is effectively in “time out” while she cares for her child. Guarding males remain in their pair bond, but multiple-mating males continue to compete for additional paternities. We do not track the individual offspring (thus do not assign a sex) until weaning, which occurs at a fixed age, *α*. Once a child is weaned (independent in our model), mothers transition to “time in” and a sex assigned with equal probability for male or female. The offspring then enter the juvenile period of their lifespan and are designated a mating strategy. For simplicity, we do not consider the dynamics of parental care and assume all juveniles are independent even though, for humans, there is a distinct period between weaning and full independence within which youngsters rely on subsidies other than their mother’s milk, so support is not provided by her alone. Males explicitly inherit the mating strategy trait and females become carriers of the trait. At time *σ*_*m*_, male juveniles become sexually mature and subsequently leave the juvenile period. Juvenile females become sexually mature at age *σ*_*f*_ (*L*), which is a function of the individual’s expected adult lifespan, *L* [2]. To focus on the core dynamics, we assume adult mortality rates are fixed, so that each juvenile and female has a lifetime mortality rate of 1*/L*, adult males have a slightly higher mortality rate of 1.09*/L* corresponding to a 9% greater chance of death. Additionally, the population is subject to an intrinsic, population-dependent death rate that affects all individuals equally. If the population surpasses a carrying capacity, *K*, an individual is randomly selected with uniform probability and removed. If the algorithm selects a female with an offspring, the dependent is also removed.

### 2.2 Fertility Window

For simplicity, we assume all females are no longer fertile after the age of 45. Thus, females are only fertile between the ages of *σ*_*f*_ (*L*) and 45. When a female experiences an end to her fertility window, she transitions to the population of postmenopausal females, *X*. If a female reaches the age of 45 while caring for a dependent, she transitions to *X*_*g*_ or *X*_*m*_, depending on the child she is caring for. All males are fertile starting at age *σ*_*m*_ and remain in this status until they reach the age of frailty and are removed from the mating pool. This age is defined as the minimum between 2*L* and 75.

### 2.3 Mating and Conception

Only females without dependants who are within their fertility window are eligible to conceive. For simplicity, we assume that these females conceive and give birth at a constant rate, *ρ*. Additionally, we assume that females have no preference for guarding or multiple-mating males, so the proportion of paternities going to guarding males is the frequency *G/*(*G* + *M*) and similarly, the proportion going to multiple-mating males is the frequency *M/*(*G* + *M*).

We also include the effect of paternity theft, defined when a multiple-mating male steals the paternity from a guarding male. The probability of this occurring, *q*, naturally depends on the proportion of multiple-maters in the population. Thus, if *q*^∗^ represents the success rate of multiple-mating thieves, then

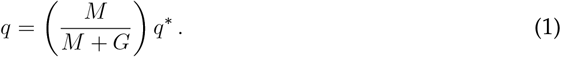

If, for example, *q*^∗^ = 0.10, then each multiple-mating male is successful at stealing a paternity 10% of the time. This means that at the population level, multiple-mating males have a success rate of ∼ 60% compared to guarding males with a success rate of ∼ 40% since without paternity theft, *q*^∗^ = 0, both multiple-mating and guarding males have an equal chance at a paternity. Like Maynard Smith’s early ESS model of parental investment [32], we represent male reproductive success probabilistically; however, our agent-based formulation differs by explicitly modelling the demographic transitions that determine mate availability.

#### 2.3.1 Trait inheritance

We assume that the multiple-mating and guarding traits are carried by the females in addition to the males. Free females are defined as either *F*^*M*^ or *F*^*G*^ which affects the trait of her offspring. Thus, if a female carrying the multiple-mating trait, *F*^*M*^, mates with a multiple-mating male, *M*, the resulting offspring will carry the multiple-mating strategy 100% of the time. On the other hand, if she mates with a guarding male, *G*, the resulting offspring will have a 50% chance of carrying the multiple-mating trait and a 50% chance of carrying the guarding trait. The same argument holds for *F*^*M*^ females. See figure S1 in the Supplementary Information, for an illustration of this process.

### 2.4 Parameter estimates

Conception times vary among great apes [24], but human data suggests an average of 4 months, so we estimate a female conception rate of *ρ* = 3 yr^−1^, corresponding to an average conception time of 1*/*3 yr, or 4 months [7].

Based on life-history regularities across primates, we assume key life-history transitions scale with respect to expected adult lifespans, *L* [2]. For mean adult lifespans, we use expected lifespans from age 15 (*e*_15_), which are 23.1 ≈ 23 for chimpanzees and 37.7 ≈ 38 for humans [36, Table 1]. Additionally, for our scaling, we assume females become sexually mature at age *L/*2.5 + 2 [22]. With this assumption, the age of female sexual maturity for chimpanzees and humans are 22*/*2.5 + 2 = 10.8 and 38*/*2.5 + 2 = 17.2, which fall near the ages of first birth in Robson *et al*. [39, Table 2.1]. Then, we assume that females reproduce up to age 45, which is close to the end of female fertility observed in both chimpanzees and humans [12]. Parameter estimates are listed in Table 1.

**Table 1.** Model parameters ane estimates.

| Parameter | Description | Estimate |
| --- | --- | --- |
| $\rho$ | Female conception rate ($\text{yr}^{-1}$) | 3 |
| $\delta_a$ | Adult/Juvenile death rate ($\text{yr}^{-1}$) | $1/L$ |
| $\delta_m$ | Adult male death rate ($\text{yr}^{-1}$) | $1.09/L$ |
| $\delta_d$ | Dependant death rate ($\text{yr}^{-1}$) | 0.08 |
| $q$ | Paternity uncertainty | 1 |
| $q^*$ | Multiple mating theft success rate | $0 \leq q^* \leq 1$ |
| $L$ | Expected adult lifespan (yr) | 23 (chimp), 38 (human) |
| $K$ | Population carrying capacity | Variable |
| $\alpha$ | Age of offspring independence from mother | Variable |
| $\chi$ | Pair-bond break-up rate | Variable |
| $\sigma_m$ | Male sexual maturity age | 15 |
| $\sigma_f(L)$ | Female sexual maturity age | $L/(2.5) + 2$ |

## 3 Results

Throughout our analysis, we investigate the parameter space corresponding to the interbirth interval length and mean adult lifespan. For each combination of these two parameters, the number of males who employ either strategy is recorded. The strategy that results in the largest population of males at equilibrium dominates and the corresponding colour is filled in. If the population goes extinct, no colour is entered. For all of those that do not go extinct, except for a few at the boundary of the two regions, only one population survives at equilibrium (see Supplementary Information, Figure S2). On each grid of results, points representing the average human- and chimpanzee-like populations are identified for comparison. To provide a baseline with which to compare parameters, we start with the most ideal situation where paternity uncertainty, *q*, and average pair-bond break-up rate, *χ*, are set to zero. The results are shown in Figure 2.

**Figure 2.**
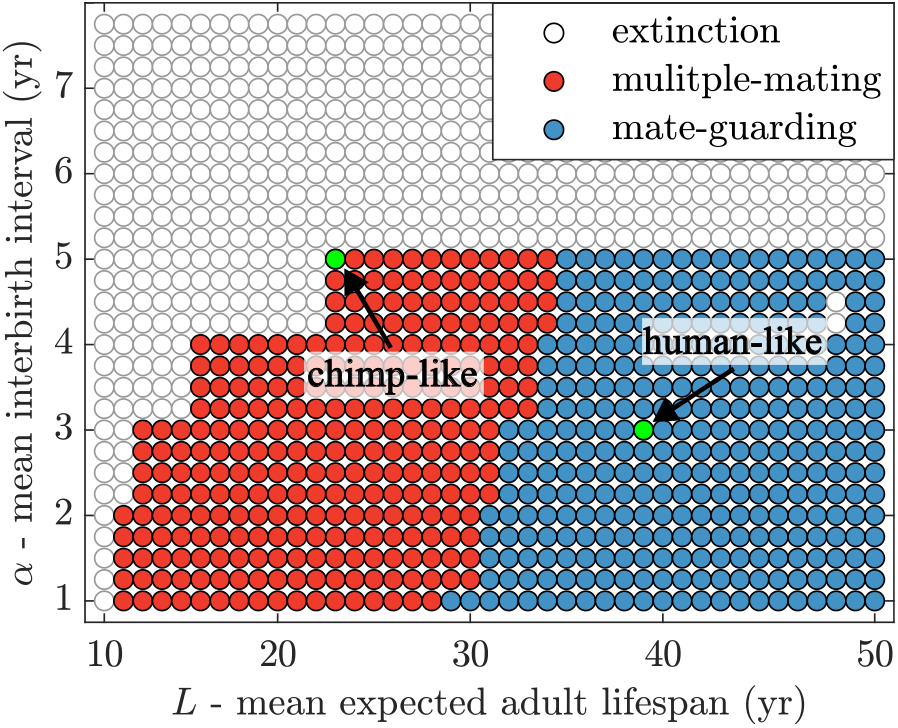
Dominant strategy when the age of independence from mothers and mean expected adult lifespan are varied. Going up the y-axis, the interbirth interval increases, making it longer before females can have their next child. Increasing x-values represent longer average adult lifespans, so the slight diagonal shift in the boundary between regions (more red at longer interbirth intervals) is the result in a trade-off between the time required for males to wait for the next paternity and expected longevity. When the interbirth interval is high, the average lifespan must be long enough to provide an advantage to guarding males. Results are shown for the idealised case in which *q* = 0 and *χ* = 0. At each point on the figure, except for a few points at the boundary between regions, the dominant strategy is the sole surviving population. Points corresponding to chimpanzee- and human-like parameters are indicated for comparison.

Note the clear distinction between chimpanzee- and human-like populations. The transition between regions in which multiple-mating and guarding dominate is driven largely by the mean expected adult lifespan, *L*. As adult lifespans expand, increasing numbers of older, still fertile males compete for paternities, but at the same time females in these ages are no longer fertile. This causes the male-biased OSR to grow in strength as *L* increases. These results suggest that guarding becomes an advantageous mating strategy as life expectancy increases, resulting in more paternity opportunities over a lifetime. The diagonal shift in the boundary between regions–more multiple mating at longer interbirth intervals–results from the trade-off between the required time for males to wait for the next paternity and their expected lifespan. If a male’s lifespan is short (male sexual maturity starts at 15), longer interbirth intervals are somewhat less advantageous for guarding males, since they may only live to experience one or two paternity opportunities. In this case, even if the competition is relatively high, there is a stronger chance of multiple opportunities in a lifetime. In the next sections, we investigate some of the underlying mechanisms behind this result and determine what happens when the idealised conditions assumed here are relaxed.

### 3.1 Male-biased sex ratios

In this section, we investigate the effects of a male-biased sex ratio on our results. We consider both the OSR and ASR. Each ratio, defined in terms of the model variables, are defined in the Supplementary Information, Table S2.

In our previous work [37], we showed that these sex ratios help to predict the transition between regions where one strategy dominates. Here, we focus on two life history equilibria associated with chimpanzee- and human-like populations where grandmothers’ subsidies allow mothers to have their next babies sooner. We start by illustrating how these sex ratios affect the dynamics of the system within the full parameter space from Figure 2. To do this, we evaluate both the ASR and OSR at each point and plot the regions where they are constant, shown in Fig. 3. Note that both ratios broadly predict the shift between regions where one strategy dominates. Generally, a higher ratio corresponds to the region where guarding dominates and a lower ratio where multiple mating is more successful. The regions of constant OSR in Figure 3(a) show far more variation than regions of constant ASR in Figure 3(b) since the OSR takes into account those who are able to conceive at a given point in time. Thus, the length of the interbirth interval has a greater effect on the sex ratio, which suggests that the OSR is more sensitive to the underlying dynamics of this system.

**Figure 3.**
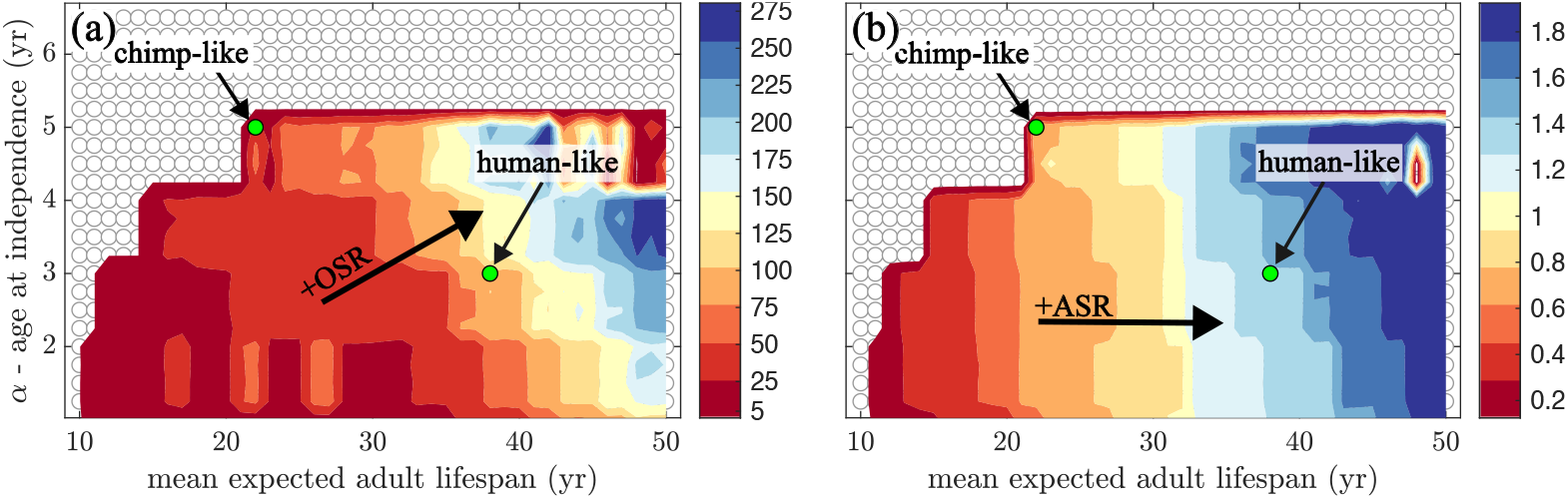
The effect of OSR and ASR on the dominant strategy in parameter space. (a) Contour regions of constant OSR generally increase from left to right and slightly up to the right for higher *α*. The arrow indicates increasing OSR. (b) Contour regions of constant ASR are more vertical, increasing from left to right for higher mean adult longevity. The arrow indicates increasing ASR. Parameters and initial conditions are the same as in Figure 2.

### 3.2 Population-level paternities

In this section, we investigate the total paternities at the population level associated with each mating strategy and continue to focus on the length of the interbirth interval and average adult longevity. Figure 4 illustrates the effect of these two parameters on the male mating strategy and their connection to the OSR. Figure 4(a) shows the OSR at equilibrium for two fixed interbirth interval lengths and varying *L*. Note that in both cases, a sudden change in the slope of the total OSR (knee of the curve) roughly corresponds to the shift in the successful mating strategy at equilibrium (red arrows). Figures 4(c) and 4(d) show the collective total number of paternities won by males using one of the two mating strategies as a function of *L* for *α* = 5 and *α* = 3 respectively. In both subfigures, red points indicate a multiple-mating strategy and blue points a guarding strategy (shades of these colours represent different lengths of the interbirth interval).

**Figure 4.**
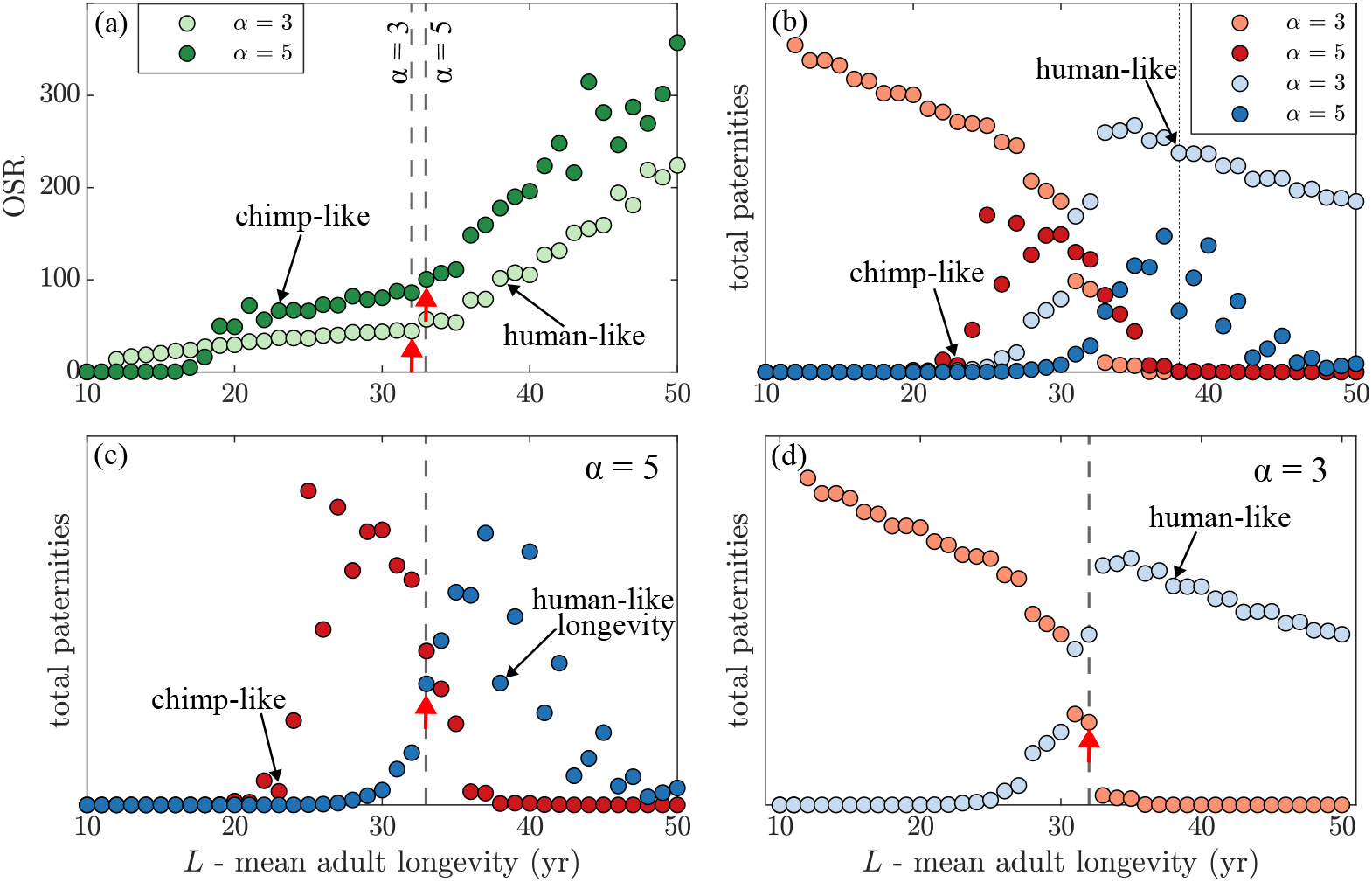
Population shift in total paternities predicted by the OSR. (a) A comparision of the OSR for a chimp-like interbirth interval of 5 years (*α* = 5) to a human-like interbirth interval of 3 years (*α* = 3). The red arrows indicate the approximate location of the knee of the curve (sudden change in slope) for both sets of data. In panels (b)-(d), the total collective paternities won by each type of male using one of the two mating strategies is shown as a function of mean adult longevity, *L*, for various fixed lengths of the interbirth interval, *α*. (c) The arrow corresponding to the chimp-like populations lines up with the shift in strategy for that fixed interbirth interval length, *α* = 5. (d) Similarly, the arrow corresponding to the human-like population lines up with the shift in strategy for total paternities for that fixed interbirth interval, *α* = 3. A comparison of both human and chimp-like populations committed to either mating strategy is shown in (b). Note that humans obtained an added benefit from reducing the interbirth interval to 3 from the ancestral chimp-like interbirth interval length of 5 years. In all cases, the age of male sexual maturity is fixed, *σ*_*m*_ = 15.

Figure 4(b) combines all of the total paternity values for both chimpanzee- and human-like interbirth interval lengths along with both mating strategies for comparison. The model predicts that humans get an added benefit by shortening their interbirth interval (dark blue to light blue). Also note that even if the interbirth interval remains at 5 years (dark blue), the human-like point (labelled human-like longevity) still produces more paternities. This adds αsupport to the grandmother hypothesis according to which extended lifespans evolved without selecting for continued female fertility at older ages [9, 10, 22, 23]. This, in turn, created conditions for a shift toward guarding. Grandmother provisioning, which allowed the inter-birth interval to drop, provided the added benefit necessary for this strategy to endure.

Results shown in figure 4 suggest that one of the primary mechanisms for the shift in strategy is a higher OSR, which is a measure of the competition facing males for each paternity opportunity. Observing the total population-level paternities won by each class of male (multiple mating and guarding), it is clear that this matches the regions shown in Fig. 2.

### 3.3 Relative paternity advantage

In this section, we introduce a ratio to measure the relative advantage of each mating strategy, which allows us to visualise the underlying dynamics associated with the transition between regions. The maximum-normalised relative difference between guarding and multiple-mating strategies is defined by the ratio

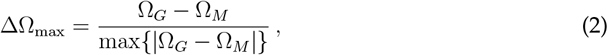

where Ω_*M*_ is the total number of paternities leading to multiple-mating offspring and Ω_*G*_ the total number of guarding offspring for the full population. In figure 5, contour lines of constant ΔΩ_max_ relative to the guarding strategy are shown (positive values indicate an advantage for guarding and negative for multiple mating).

**Figure 5.**
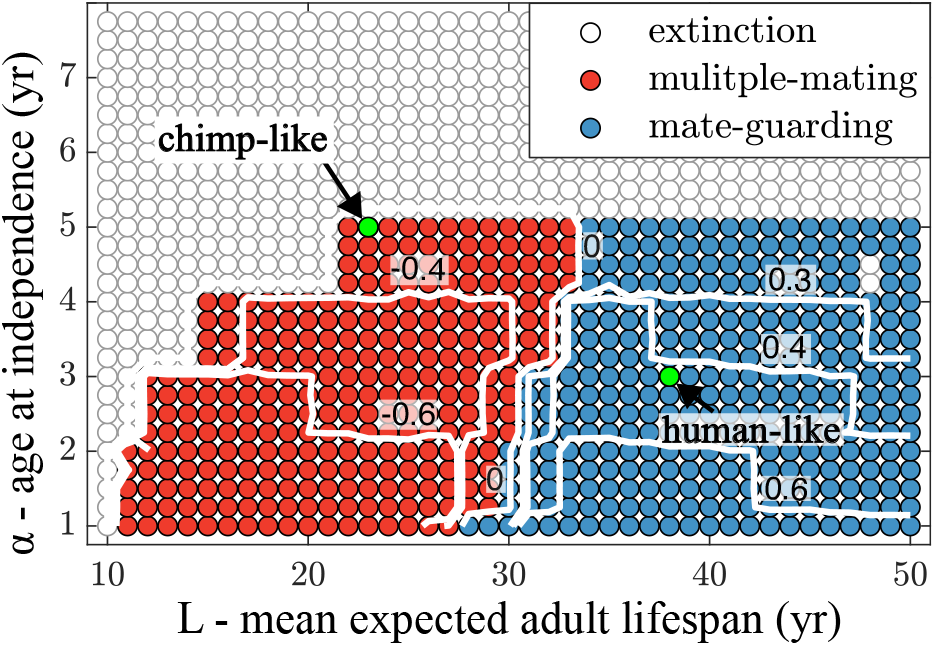
Maximum-normalised difference between guarding and multiple mating strategies, ΔΩ_max_. The relative advantage of the guarding strategy is shown by constant contour lines. Positive values indicate an advantage for guarding and negative values an advantage for multiple mating. Note that the line of zero difference approximates the shift in strategy. both chimpanzee- and human-like populations are included for comparison (green dots).

Notice that the zero line occurs at the boundary of the two regions, which clearly defines the shift in successful strategy. The relative advantage of the multiple-mating strategy increases with higher values of *α*, supporting the hypothesis that the observed diagonal shift in the boundary along the vertical axis arises from a trade-off between the time required for males to secure subsequent mating opportunities and their expected longevity.

### 3.4 Paternity uncertainty and pair bond duration

In this section, we explore the effects of paternity uncertainty and pair bond duration on the dynamics of the system. In each case, we compare the corresponding chimpanzee- and human-like populations to determine how these parameters affect the transition between regions.

Figure 6 shows the effect of increasing paternity uncertainty, *q*, measured relative to the theft success rate per multiple-mating thief, *q*^∗^. Greater paternity uncertainty makes it less advantageous to invest effort into mate guarding, thus with a higher success rate, the guarding region shrinks. The model predicts that the success rate must be no more than 10% for human-like populations to be within the mate guarding region. When the theft success rate is as high as 15%, human-like populations are no longer in the guarding region.

**Figure 6.**
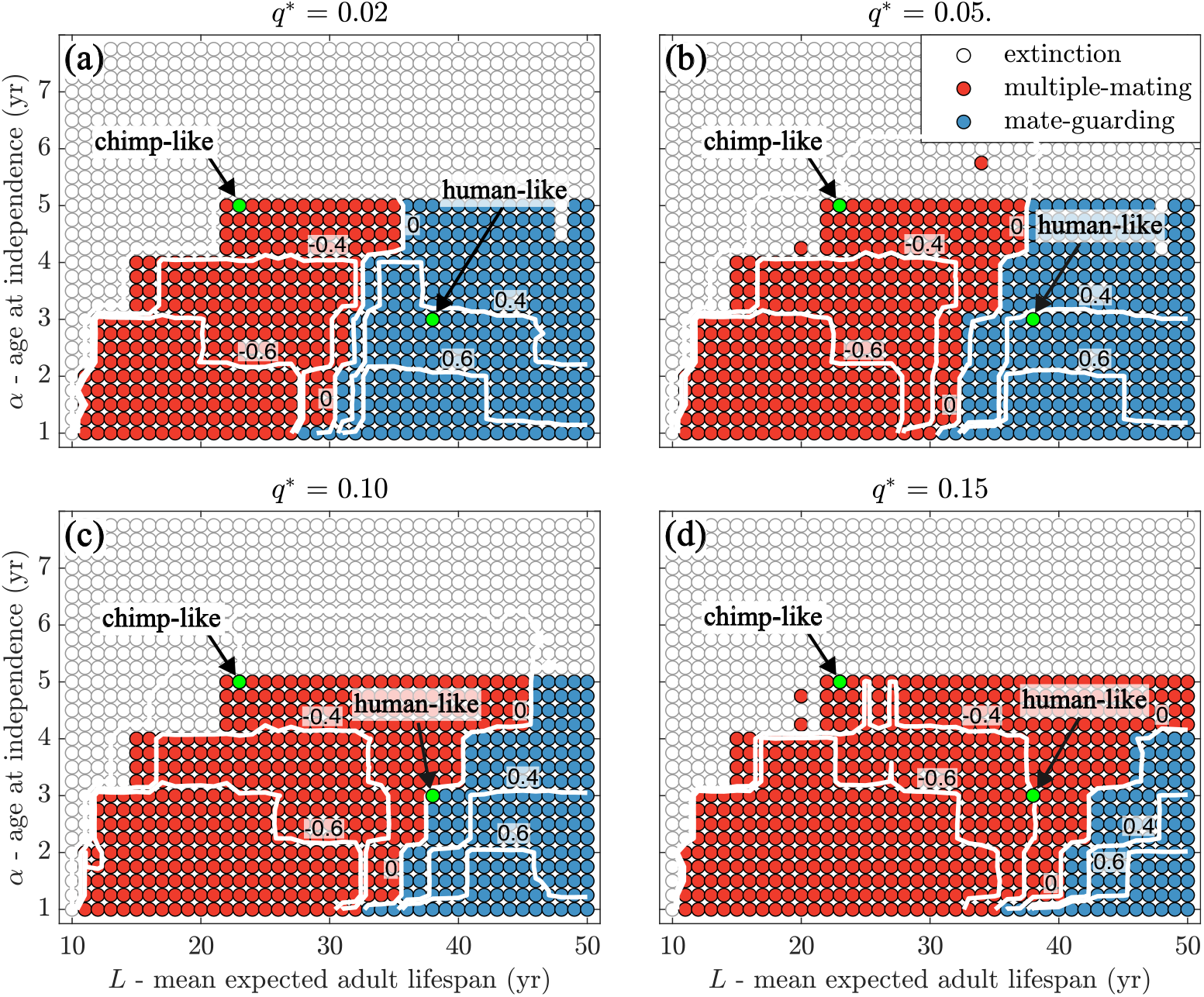
Dominant strategy for different levels of paternity uncertainty. Contour lines show constant relative advantage of guarding, ΔΩ_max_. Negative values mean an advantage for multiple mating and positive values an advantage for guarding. Results are shown for increasing paternity uncertainty when the theft success rate per multiple-mating thief is (a) 2%, 5%, (c) 10%, (d) 15%. As theft success increases, the guarding region shrinks. When the success rate is 10%, the human-like population is right on the border between regions and by the time there is 15% uncertainty, human-like populations are no longer in the guarding region. Recall that these are equilibrium populations achieved over thousands of years and that each multiple-mating male not only gets 100% of their usual paternities, but also erodes the guarding benefit by a fixed probability (guarding males stay faithful and wait for the next paternity). Thus, results illustrate the robust nature of guarding. Contour lines of zero advantage align well with the boundary between regions. In each case, the age at sexual maturity for males is *σ*_*m*_ = 15.

Recall that these values represent equilibrium populations after running the model for over a thousand years. Thus, each multiple-mating male thief not only secures 100% of his own paternities, but also captures 10% of those from guarding males. Consequently, at the population level, multiple-mating males obtain about 60% of all paternities, while guarding males are left with only 40%. This highlights the robustness of the guarding strategy for human-like populations, which can maintain reproductive success even under persistent theft.

In Figure 7, we explore how the average pair bond duration affects the dominant strategy. As pairs last longer, the mate-guarding region increases in size, indicating that it is a more favourable strategy for a broader range of parameters. This occurs since it is less advantageous to invest in mate guarding if the average pair bond is short-lived. In figure 7(a), the mean pair bond lasts only one year, which is equivalent to multiple mating. However, as the mean pair bond length increases to 5, 15 and 30 years, as shown in figures 7(b), (c), and(d), the guarding region increases. The model predicts pair bonds must last 15 years on average for human-like populations to benefit from guarding.

**Figure 7.**
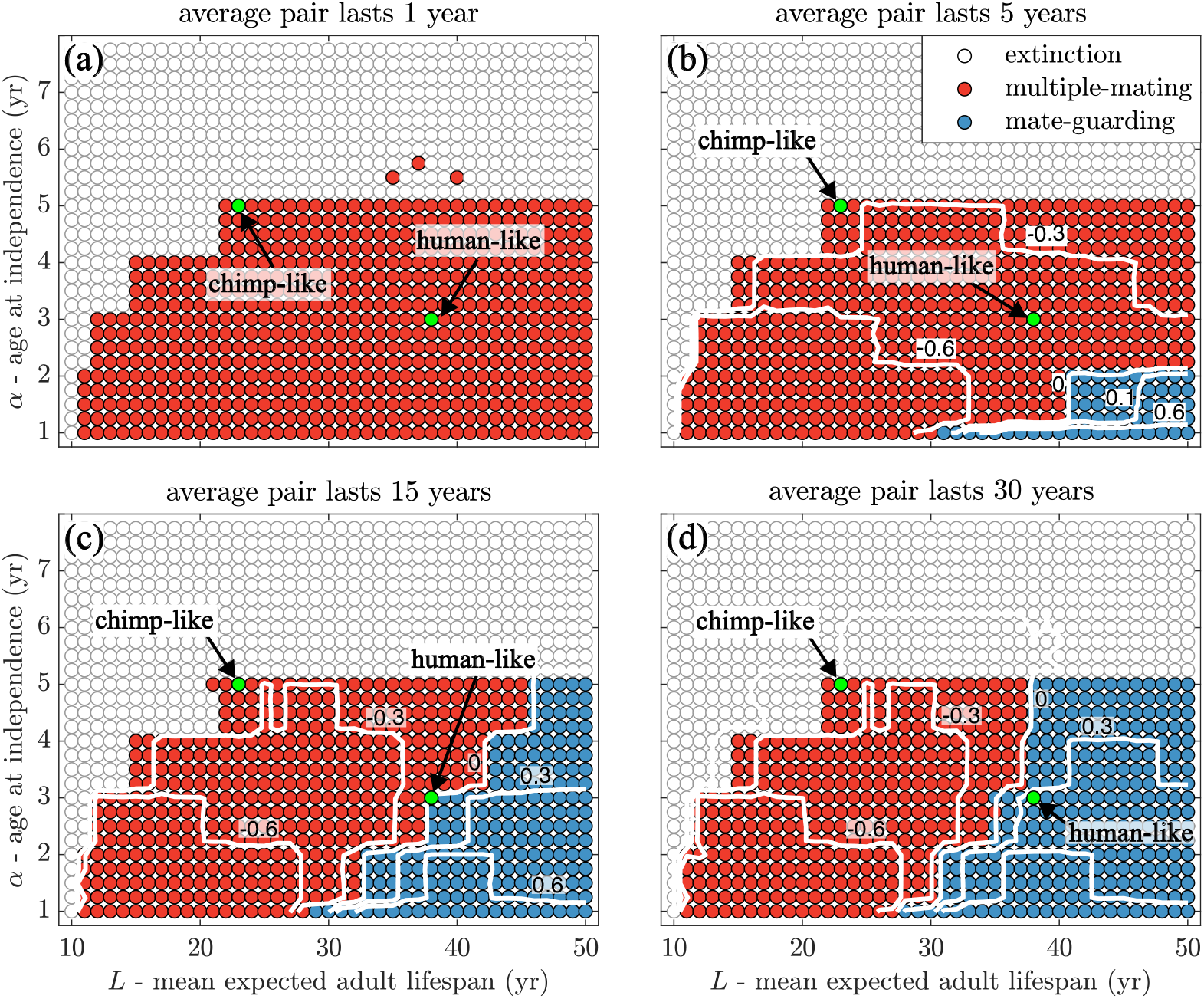
Dominant strategy for different pair-bond duration lengths. Contour lines show constant relative advantage of guarding, ΔΩ_max_. Negative values mean an advantage for multiple mating and positive values an advantage for guarding. Results are shown for increasing average pair-bond duration of (a) 1 year, (b) 5 years, (c) 15 years, and (d) 30 years. If pair bonds only last 1 year, they are equivalent to multiple mating. However, as the average length of the pair bond increases, the blue region also increases in size. When the average pair lasts 15 years, the human-like population is right on the border between regions and by the time the bonds last 30 years, human-like populations are well within the guarding region. Contour lines of zero advantage align well with the boundary between regions. In each case, the age at sexual maturity for males is *σ*_*m*_ = 15

Based on the results, varying these two parameters significantly affects the dynamics of the system. In particular, more uncertainty due to paternity theft leads to a larger region of multiple mating and longer pair bonds leads to a larger region where guarding dominates. Figure 8 illustrates how these parameters work together for human-like populations (*L* = 38, *α* = 3). The full results are shown in figure 8(a), with the ΔΩ_max_ = 0 contour line super-imposed which predicts the boundary between regions. Moving up the y-axis, the success rate per multiple-mating thief, *q*^∗^, increases, raising paternity uncertainty. Increasing values along the x-axis indicate lower mean breakup rates, *χ*, which generally improve guarding performance. However, greater paternity-theft success erodes the effectiveness of guarding, causing the guarding region to shrink from the bottom to the top of the figure. These results should be interpreted in the context of chimpanzee-like guarding relationships, which differ from the more complex, persistent, and emotionally rich pair bonds of human hunter-gatherers. Our aim is to trace the evolutionary origins of human pair bonds, approaching the question from the chimpanzee end of the spectrum in order to better understand how this behaviour may have emerged. Figure 8(b) shows regions of constant relative advantage of guarding, illustrating the strength of that strategy away from the border between regions. The results show that the transition between regions is sharp with one strategy quickly replacing the other.

**Figure 8.**
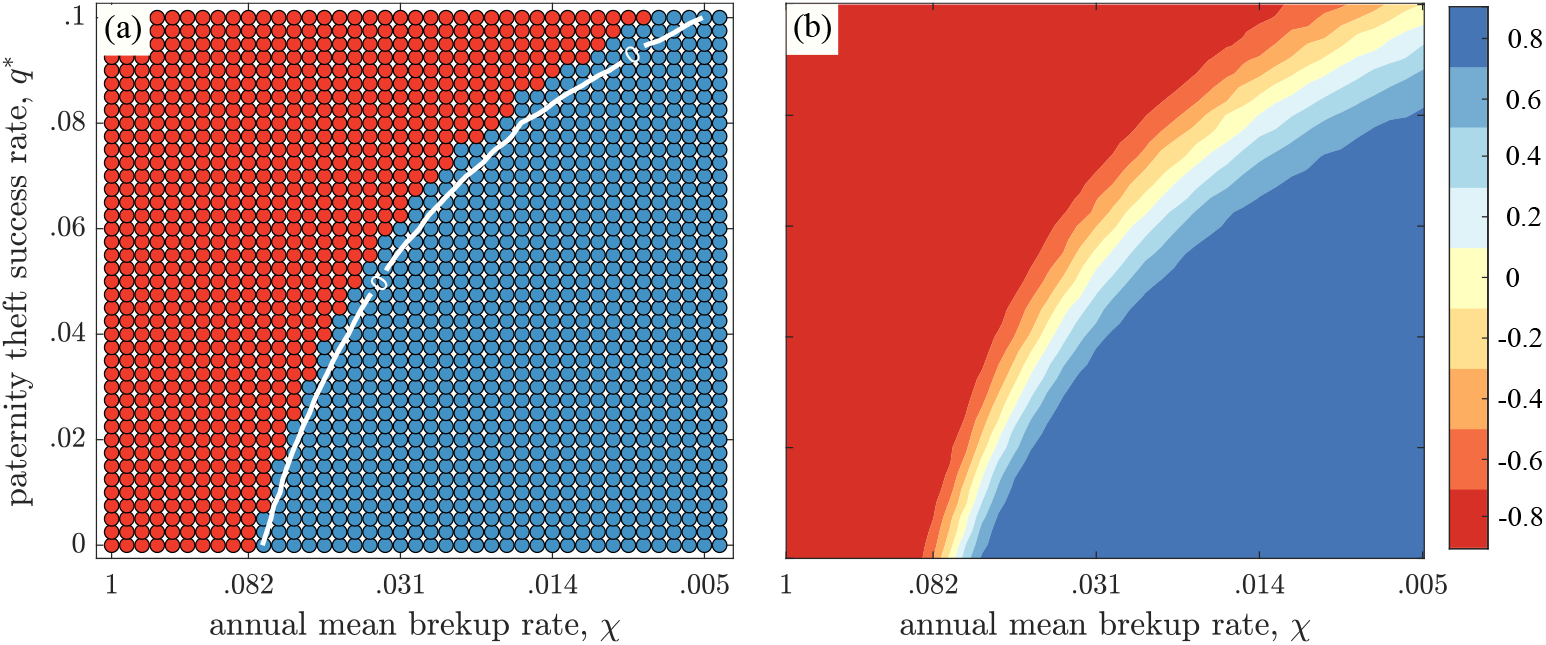
Dominant strategy when both paternity-theft success *q*, and mean breakup rate, *χ*, vary for human-like populations (*L* = 38, *α* = 3). (a) The dominant strategy showing the line of zero constant ΔΩ_max_ at the boundary between regions. Increasing values on the y-axis represent higher paternity-theft success per multiple-mating thief, while larger x-axis values correspond to a lower annual mean breakup rate, *χ*. Higher paternity uncertainty expands the multiple-mating region. Conversely, lower mean breakup rates result in a larger guarding region. The curved boundary arises because guarding erodes rapidly as thieves become more successful. Results should be viewed through the lens of chimp-like guarding relationships (our starting point for tracing the roots of human pair bonds) which are unlike the more complex, persistent and emotionally distinct human marriages. (b) Contour regions showing the relative advantage of guarding over multiple-mating highlight a sharp transition between regions, indicating that the shift from guarding to multiple mating is rapid, not gradual.

## 4 Conclusion

We have developed an agent-based, age-structured model to investigate the factors affecting the evolution of male mating strategies in the human lineage. Our goal in constructing such a model is to understand the key parameter combinations that contribute to the shift from a multiple-mating strategy in chimpanzee-like populations to a mate-guarding strategy in human-like populations. Although there are multiple factors that contribute both directly and indirectly to the establishment of mate guarding, we deliberately take a simplifying approach to avoid any choice regarding the functional relationship between them. In particular, we focused on the two most direct factors—the interbirth interval and expected adult longevity (which most importantly includes postmenopausal longevity)—and did not take into account male-male or male-female interactions. We only were concerned with the equilibrium population as they relate to mean fertility and mortality rates.

The results indicate that the shift from multiple-mating to guarding as a successful male mating strategy is linked with a higher male-biased sex ratio (both ASR and OSR) which corresponds to a longer interbirth interval and mean adult longevity. It is, however, the OSR that most closely aligns with the underlying mechanism for this transition, which measures the strength of competition for each available female. We found a correlation when we directly compared the OSR with the population-level shifts in total paternities achieved by males using a given mating strategy. The OSR reaches a point of sudden change in the slope (knee of the curve) at approximately the same point where the total paternities secured by the multiple-mating strategy diverge from those achieved through mate guarding.

The grandmother hypothesis posits that support from grandmothers to their daughters’ children shortened interbirth intervals, driving the evolution of extended lifespan without selecting for continued female fertility at older ages [9, 10, 22, 23]. In our model, post-fertile longevity increases competition for fertile females, making mate guarding the dominant strategy [3]. Our results suggest that as ancestral grandmothers’ productivity propelled the evolution of increased postmenopausal longevity by subsidising the dependants of still-fertile females, effects on male mating strategies were doubly consequential. The increased longevity included older still-fertile males which pushed the sex ratio of the fertile pool (the ASR) to become male-biased while the shorter birth intervals also expanded the paternity opportunities per male (the OSR).

While our analysis of this model yields fascinating insights, it also highlights several limitations and unresolved questions. One of the key distinctions between humans and chimpanzees (and mammals in general) is the degree to which weaned dependants become independent feeders. Chimpanzees and other mammals begin foraging while still nursing, so independence from the mother coincides with true self-sufficiency. In contrast, human offspring remain highly dependent after weaning. Consequently, while our simulations do not explicitly model the direct subsides provided by post-fertile females, we approximate their effects through shorter inter-birth intervals, an advantage made possible in humans by such support. Investigating the dynamics of this care would be a valuable extension of the model. Also, relying on direct patrilineal transmission of strategy is not realistic; it would be valuable to explore how individuals actually inherit or adjust their strategic tendencies in response to social and cultural factors over the course of their lives. Additionally, how does male-male competition affect the outcome? The average age of first reproduction for humans is female-biased (17 for females and 21 for males [1, 31]). Even though the younger males are in the fertile ages, they do not get paternities because the older males out-compete them. This is a divergence from chimpanzee behaviour where adolescent males actually get paternities on young females as the older males (who prefer older females [35]) do not prevent them [33]. Furthermore, there may be other techniques males employ that affect the winning strategy at equilibrium. Guarding males may aid their offspring through provision of care and protection from infanticide. Thus, there are important dynamics not captured by this model.

Also, we view the formation of pair bonds through the lens of chimpanzee guarding. This is a simplifying assumption that does not take into account the complexities of human marriage. It has recently been suggested that cuckoldry is more common in human hunter gatherer marriages than previously realised [41]. Genetic testing within communities of Himba pastoralists living in Namibia, revealed that nearly 50% of children born to married couples are fathered by someone other than the husband [42]. Allowing guarding males to be unfaithful in our model would certainly increase the size of the resulting guarding regions we observed. Guarding males would not only benefit from a guarded female but also could occasionally benefit from additional paternities through other unguarded females when his partner is unavailable for conception. However, this brings to light another unresolved question. If cuckoldry is common in humans, why are marriages so prevalent? In modern human hunter gatherer societies, such as the Hadza of northern Tanzania, men who are known to be expert hunters re-marry faster after a breakup and claim younger women with each marriage [1].

Finally, where did the practice of guarding originate? Scholars have long debated its evolutionary roots, suggesting that a mix of ecological pressures and cultural developments played a role. The leading hypothesis proposes that guarding arose alongside cooperative parenting, offering collective benefits [5,18,19, 21, 29, 45]. One of the first models to explore an alternative hypothesis [11] found that under a wide range of conditions, guarding could become the favoured strategy. Our own model lends further credence to the idea that human pair-bonding co-evolved with increasing advantages for mate guarding—a dynamic driven in part by the emergence of a “grandmothering” life history [3]. Nonetheless, many facets of this evolutionary pathway remain to be fully understood.

## Data Availability

The authors confirm that the data supporting the findings of this study are included in the article. All relevant Matlab code and data for the analyses and results presented are available at https://github.com/mcnitschke/ABMmalemating

## CRediT authorship contribution statement

**Matthew C. Nitschke:** Conceptualisation, Methodology, Formal analysis, Investigation, Writing – original draft, Writing – review & editing, Visualisation. **Kristen Hawkes:** Conceptualisation, Writing – review & editing. **Peter S. Kim:** Conceptualisation, Writing –review & editing.

## Acknowledgements

The authors gratefully acknowledge support for this work through the Australian Research Council Future Fellowship FT200100190 (PSK).

