## Supplementary Information for "The male-biased sex ratio in humans and its role in the transition from promiscuity to pair bonding"

#### 1 Population variables and sex ratios

In this section, we provide a complete list of all subpopulations along with the definition of both sex ratios we consider in our model. This is a model structured by sex, fertility status, and mating strategy. Both males and females carry one of two traits corresponding to a male mating strategy. Individuals are either juvenile, actively in the mating pool, or post-fertile and retired from mating. The full list of all subpopulations is provided in Table S1.

| Variable | Description |
|---|---|
| $F^M$ | Population of free females carrying the multiple-mating trait |
| $F^G$ | Population of free females carrying the guarding trait |
| $M$ | Population of multiple-mating males |
| $G$ | Population of unpaired guarding males |
| $F_m^M$ | Population of unpaired females with the multiple-mating trait caring for dependent offspring with the multiple-mating trait |
| $F_g^G$ | Population of unpaired females with the guarding trait caring for dependent offspring with the guarding trait |
| $F_m^G$ | Population of unpaired females with the guarding trait caring for dependent offspring with the multiple-mating trait |
| $F_g^M$ | Population of unpaired females with the multiple-mating trait caring for dependent offspring with the guarding trait |
| $P_g^G$ | Population of pairs of guarding males and females with the guarding trait caring for dependants with the guarding trait |
| $P^G$ | Population of pairs of guarding males and females carrying the guarding trait without dependants |
| $P_m^M$ | Population of pairs of guarding males and females with the multiple-mating trait caring for dependent offspring with the multiple-mating trait |
| $P_g^M$ | Population of pairs of guarding males and females with the multiple-mating trait caring for dependent offspring with the guarding trait |
| $P^M$ | Population of pairs of guarding males and females carrying the multiple-mating trait without dependants |
| $J_m^M$ | Population of multiple-mating male juveniles |
| $J_m^G$ | Population of guarding male juveniles |
| $J_f^M$ | Population of female juveniles carrying the multiple-mating trait |
| $J_f^G$ | Population of female juveniles carrying the guarding trait |
| $X$ | Population of post-fertile females without dependants |
| $X_m$ | Population of post-fertile females caring for dependent offspring with the multiple-mating trait |
| $X_g$ | Population of post-fertile females caring for dependent offspring with the guarding trait |

**Table S1:** Population variables. Subpopulations are grouped by sex, fertility status, and mating strategy.

The two sex ratios we consider, defined in terms of the subpopulations, are listed in Table S2. The adult sex ratio (ASR) takes into account all males and females who are free, in committed pair-bonds, or caring for dependants. The operational sex ratio (OSR) only concerns sexually competing males and females.

| Ratio | Definition |
| --- | --- |
| Adult Sex Ratio | $\frac{M + G + P_g^G + P^G + P_m^M + P_g^M + P^M}{F^M + F^G + F_m^G + F_m^M + F_g^G + F_g^M + P_g^G + P_g^M + P^G + P_m^M + P^M}$ |
| Operational Sex Ratio | $\frac{M + G}{F^M + F^G}$ |

**Table S2:** Definition of ASR and OSR with population variables as in Table S1. Note that the ASR takes into account all males and females who are free, in committed pair-bonds, or caring for dependants. The OSR only concerns sexually competing males and females. In our model, this does not include males or females in a pair as we do not allow for guarding males to have paternities outside of pair bonds.

### 2 Mating interactions

The multiple-mating and guarding traits are carried by both the males and females. The mating interactions and the approximate method by which these traits spread through mating is illustrated in figure S1.

In Figure S1(top), free females carrying the multiple-mating trait, $F^M$, productively mate at rate $\rho$. They mate with multiple-mating males, $M$, or unpaired guarding males, $G$, with probabilities $M/(M+G)$ or $G/(M+G)$. If they mate with multiple-mating males, they transition to the population $F_m^M$ of females caring for dependent offspring with the multiple-mating trait. If they mate with guarding males, they form pairs with the guarding males and together enter the population $P_m^M$ or $P_g^M$ with equal chance (thus the factor of $1/2$). Paternity theft can occur only after the offspring matures. (Females drive productive mating at rate $\rho$, so all males, $M$ and $G$, mate at rate $\rho F^M/(M+G)$, which is $\rho$ times the proportion of free females per unpaired male.) Pairs, $P^M$ of females and guarding males without dependants also productively mate at rate $\rho$ and enter population $P_m^M$ or $P_g^M$ with equal chance.

In Figure S1(bottom), free females carrying the guarding trait, $F^G$, productively mate at rate $\rho$. They mate with multiple-mating males, $M$, or unpaired guarding males, $G$, with probabilities $M/(M+G)$ or $G/(M+G)$. If they mate with multiple-mating males, they transition to population $F_g^G$ or $F_m^G$ with equal probability (thus the factor of $1/2$). If they mate with guarding males, they form pairs with the guarding males and together enter the population $P_g^G$. Pairs, $P^G$, of females and guarding males without dependants also productively mate at rate $\rho$ and enter population $P_g^G$.(Females drive productive mating at rate $\rho$, so all males, $M$ and $G$, mate at rate $\rho F^G/(M+G)$, which is $\rho$ times the proportion of free females per unpaired male.) We include the chance that paternities from paired males can be stolen with probability $q$. In our model, this can only occur after pairs are established and the first successful conception by a guarding male.

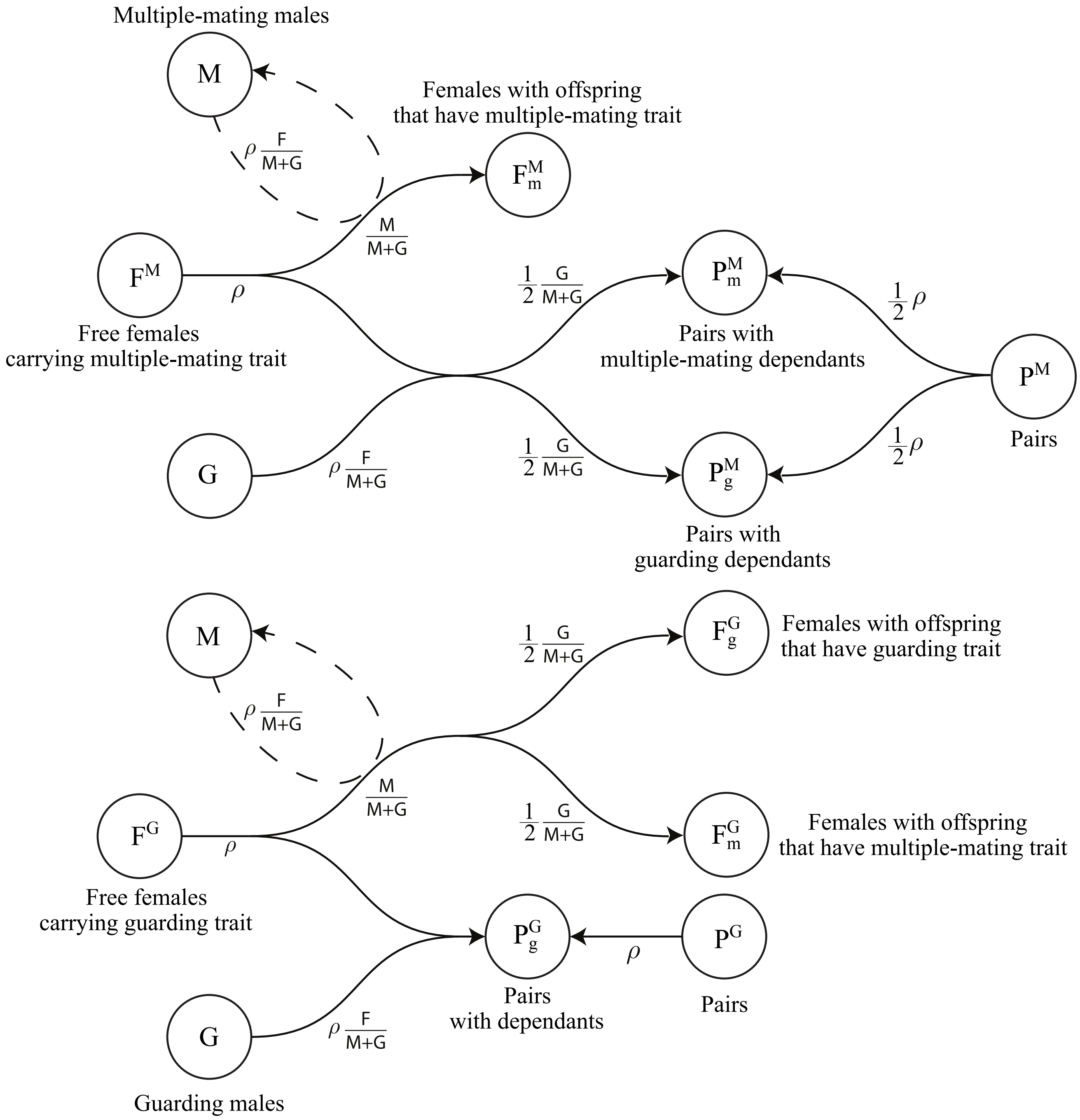


**Figure S1:** Model diagram for mating interactions. Both males and females carry the multiple-mating and guarding traits. (top) Free females carrying the multiple-mating trait, $F^M$, productively mate at rate $\rho$. They mate with multiple-mating males, $M$, or unpaired guarding males, $G$, with probabilities $M/(M+G)$ or $G/(M+G)$. If they mate with multiple-mating males, they transition to the population $F^M_m$ of females caring for dependent offspring with the multiple-mating trait. If they mate with a guarding males, they either transition to the population of pairs with multiple-mating dependants, $P^M_m$ or the population of pairs with guarding dependants, $P^M_g$. Each possibility is given an equal chance. (bottom) Free females carrying the guarding trait, $F^G$, productively mate at rate $\rho$. They also mate with multiple-mating males, $M$, or unpaired guarding males, $G$, with probabilities $M/(M+G)$ or $G/(M+G)$. If they mate with guarding males, they transition to the population $P^G_g$ of pairs with guarding dependants. If they mate with multiple-mating males, they either transition to the population of females with offspring that have the guarding trait, $F^G_g$ or the population of females with offspring that have the multiple-mating trait, $F^G_m$. Each possibility is given an equal chance.

### 3 Further results

In this section, we provide further details regarding the baseline results. First, we provide a contour plot showing the overlap between regions illustrated in Figure 2 of the main text. The results show that the transition between regions is rather abrupt. Recall that these values represent the long-term equilibrium populations that remain after more than 1000 years. Second, we illustrate two extreme cases that do not fit the assumptions of the model: females experience a loss of fertility during their lifetime and guarding males re-enter the mating pool when the female they are paired with loses fertility. For completion, we show how these key assumptions affect the results.

#### 3.1 Region overlap

A contour plot of relative reproductive success, $R_i$, for males employing the mate-guarding and multiple-mating strategies is shown in Figure S2. The $x$-axis represents mean longevity (expected male lifetime), and the $y$-axis represents the interbirth interval length, the average time between successive reproductive events. Each contour line connects combinations of $\alpha$ and $L$ that yield equivalent total population-level reproductive output, normalised by the maximum paternity count achieved across all strategies ($R_i = \Omega_i / \max_j \Omega_j$), where $i \in \{M, G\}$ for multiple mating and guarding (recall that $\Omega_M$ and $\Omega_G$ are the total number of paternities leading to multiple-mating or guarding offspring). Regions of high relative success ($R_i \approx 1$) indicate conditions under which a given strategy approaches the reproductive performance of the most successful alternative, whereas lower values correspond to reduced fitness relative to that optimum. This highlights the relationship between the different strategies as success changes from one region to the next. Note that except for a small region of overlap, a single strategy survives.

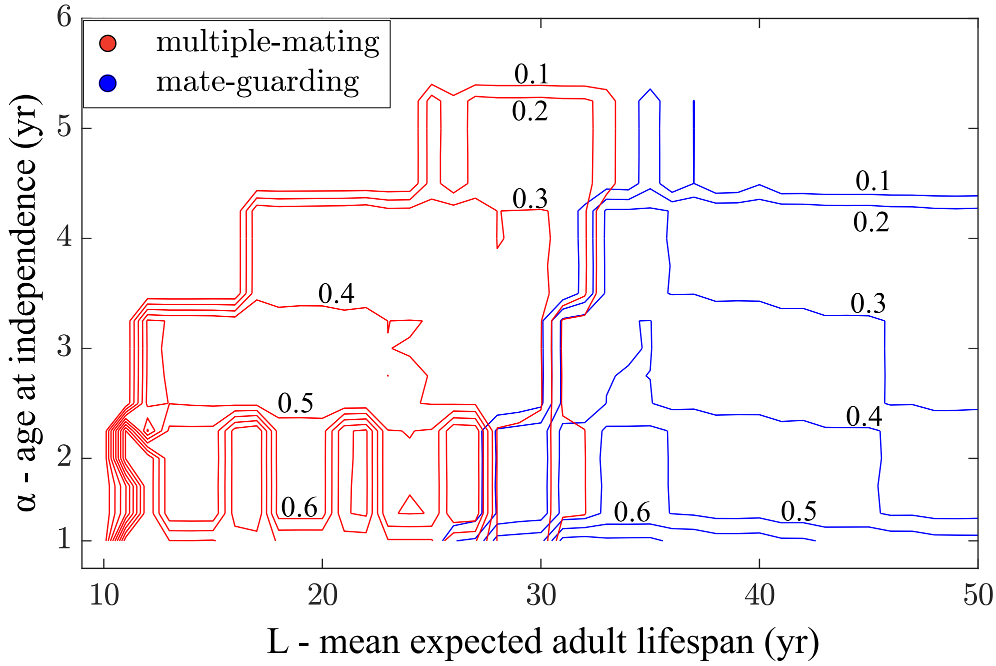


**Figure S2:** Relative reproductive success of each mating strategy. Contour plot of relative reproductive success for two competing male mating strategies, mate-guarding and multiple-mating. Each contour line represents combinations of $\alpha$ and $L$ that yield equal total population-level reproductive success, normalised to the most successful strategy (relative reproductive success = 1). Values closer to 1 indicate conditions under which a given strategy achieves comparable paternity output to the dominant strategy. The plot illustrates the fitness landscape across which shifts in ecological or demographic parameters may alter the relative advantage of mate-guarding versus multiple-mating.

#### 3.2 Extreme cases

In this section, we investigate several of the important underlying model assumptions by considering extreme cases. In particular, we determine the results when females remain fertile for as long as men and when male guarders leave the mating pool when their female pair-mate loses fertility.

In order to isolate the effects of the fertility window and how male and female sexual maturity age play a role, we consider the model when there is no end to female fertility, shown in Figure S3(a). When the fertility window is not bounded by old age, we observe that later ages at offspring independence are allowed without causing extinction. Multiple mating is far more prevalent since there are always an abundance of females available for more overall paternity opportunities.

In our model, we assumed paired males break up and re-enter the mating pool when their partners lose fertility. This assumption is also incorporated in many other models [1, 3, 4]; however, the alternative assumption that males remain paired and hence cease reproducing when their partners lose fertility endures in some studies [2, 5] and thus is worth exploring. In Figure S3(b), a grid of points in parameter space is generated for this case where paired males effectively become infertile at the same time as their partners. This illustrates that under these conditions, guarding only appears for extreme values of $L$ and $\alpha$. In this scenario, both chimpanzee- and human-like populations remain well within the multiple mating region.

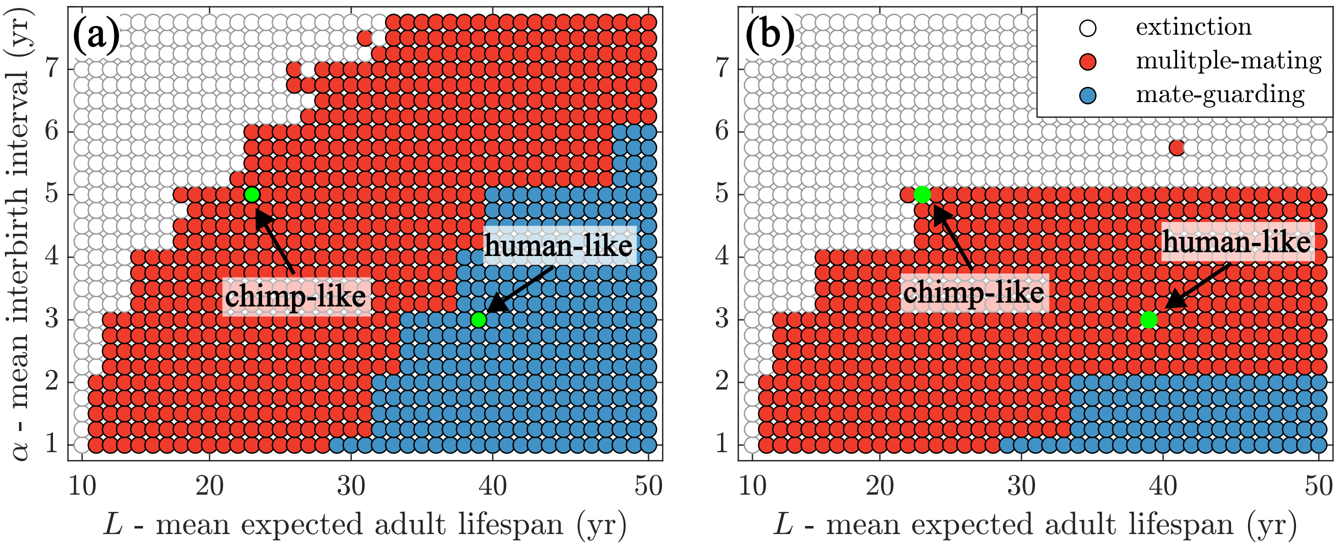


**Figure S3:** Extreme model assumptions. (a) Females do not lose fertility and live as long as males. Under this extreme condition, there is less extinction as females have a much longer fertility window and the population does not crash when the interbirth interval is extreme. (b) Guarding males remain out of the mating pool when their partner is no longer fertile. In this case, the guarding region is much smaller with human-like populations well within the multiple mating region. Thus, it's important that males with the guarding trait re-enter the mating pool, otherwise there is less advantage to this strategy.
